# Neural spiking for causal inference

**DOI:** 10.1101/253351

**Authors:** Benjamin James Lansdell, Konrad Paul Kording

## Abstract

When a neuron is driven beyond its threshold it spikes, and the fact that it does not communicate its continuous membrane potential is usually seen as a computational liability. Here we show that this spiking mechanism allows neurons to produce an unbiased estimate of their causal influence, and a way of approximating gradient descent learning. Importantly, neither activity of upstream neurons, which act as confounders, nor downstream non-linearities bias the results. By introducing a local discontinuity with respect to their input drive, we show how spiking enables neurons to solve causal estimation and learning problems.

## Introduction

Most nervous systems communicate and process information with spiking neural networks. Yet machine learning mostly uses artificial neural networks with continuous activities. Computationally, despite a lot of recent progress [58, 71, 8, 9, 52], it remains challenging to create spiking neural networks that perform comparably to artificial networks. Instead, spiking is generally seen as a disadvantage – it is difficult to propagate gradients through a discontinuity. This disparity between spiking and artificial networks raises the question, what are the computational benefits of spiking?

There are, of course, pragmatic reasons for spiking: spiking may be more energy efficient [1, 70], spiking allows for reliable transmission over long distances [59], and spike timing codes may allow for more transmission bandwidth [72]. Yet, despite these ideas, we may still wonder if there are computational benefits of spikes that balance the apparent disparity in the abilities of spiking and artificial networks.

A key computational problem in both biological and artificial settings is the credit assignment problem. When performance is sub-optimal, the brain needs to decide which activities or weights should be different. Credit assignment is fundamentally a causal estimation problem – which neurons are responsible for the bad performance, and not just correlated with bad performance? Solving such problems is difficult because of confounding: if a neuron of interest was active during bad performance it could be that it was responsible, or it could be that another neuron whose activity is correlated with the neuron of interest was responsible. In general, confounding happens if a variable affects both another variable of interest and the performance. Even when a fixed stimulus is presented repeatedly, neurons exhibit complicated correlation structures [5, 15, 37, 7, 78] which confounds a neuron’s estimate of its causal effect. This prompts us to ask how neurons can solve causal estimation problems.

The gold-standard approach to causal inference is randomized perturbation. If a neuron occasionally adds an extra spike (or removes one), it could readily estimate its causal effect by correlating the extra spikes with performance. Such perturbations come at a cost, since the noise can degrade performance. This class of learning methods has been extensively explored [11, 20, 21, 44, 49, 77, 64]. However the learning speed of a network of neurons using random perturbations scales poorly with the number of neurons *N* (as *𝒪* (1*/N*) [64, 20, 60]). Further, in general it is not clear how a neuron may know its own noise level (though see the birdsong learning literature for one plausible case [20], or other proposals about how it may be approximated in some cases [44, 30]). Thus we may wonder if neurons estimate their causal effect without random perturbations.

How could neurons estimate their causal effect? Over a short time window, a neuron either does or does not spike. Comparing the average reward when the neuron spikes versus does not spike gives a confounded estimate of the neuron’s effect. Because neurons are correlated, spiking is associated with a different network state than not-spiking. This difference in network state that may account for an observed difference in reward, not specifically the neuron’s activity. Simple correlations will give wrong causal estimates.

However, the story is different when comparing the average reward in times when the neuron *barely* spikes versus when it *almost* spikes. The difference in the state of the network in the barely spikes versus almost spikes case is negligible, the only difference is the fact that in one case the neuron spiked and in the other case the neuron did not. Any difference in observed reward can therefore *only* be attributed to the neuron’s activity. The precise mathematical reason is that the drive satisfies the back-door criterion relative to the spike-reward relation [56]. In this way the spiking discontinuity may allow neurons to estimate their causal effect.

Here we propose the spiking discontinuity is used by a neuron to efficiently estimate its causal effect. Once a neuron can estimate its causal effect, it can use this knowledge to calculate gradients and adjust its synaptic strengths. We show that this idea suggests a learning rule that allows a network of neurons to learn to maximize reward. We demonstrate the rule in simple models. The discontinuity-based method provides a novel and plausible account of how neurons learn their causal effect.

## Results

### A neuron’s causal effect

In causal inference, the causal effect can be understood as the expected difference in an outcome *R* when a treatment *H* is randomly, independently, assigned – a primary quantity of interest in a randomized control trial. This definition can be made precise in the context of a causal graphical model [56] (Figure 1A, see Methods; or with the potential outcomes framework [3]). Notationally, the causal effect can be defined as:

**Figure 1:**
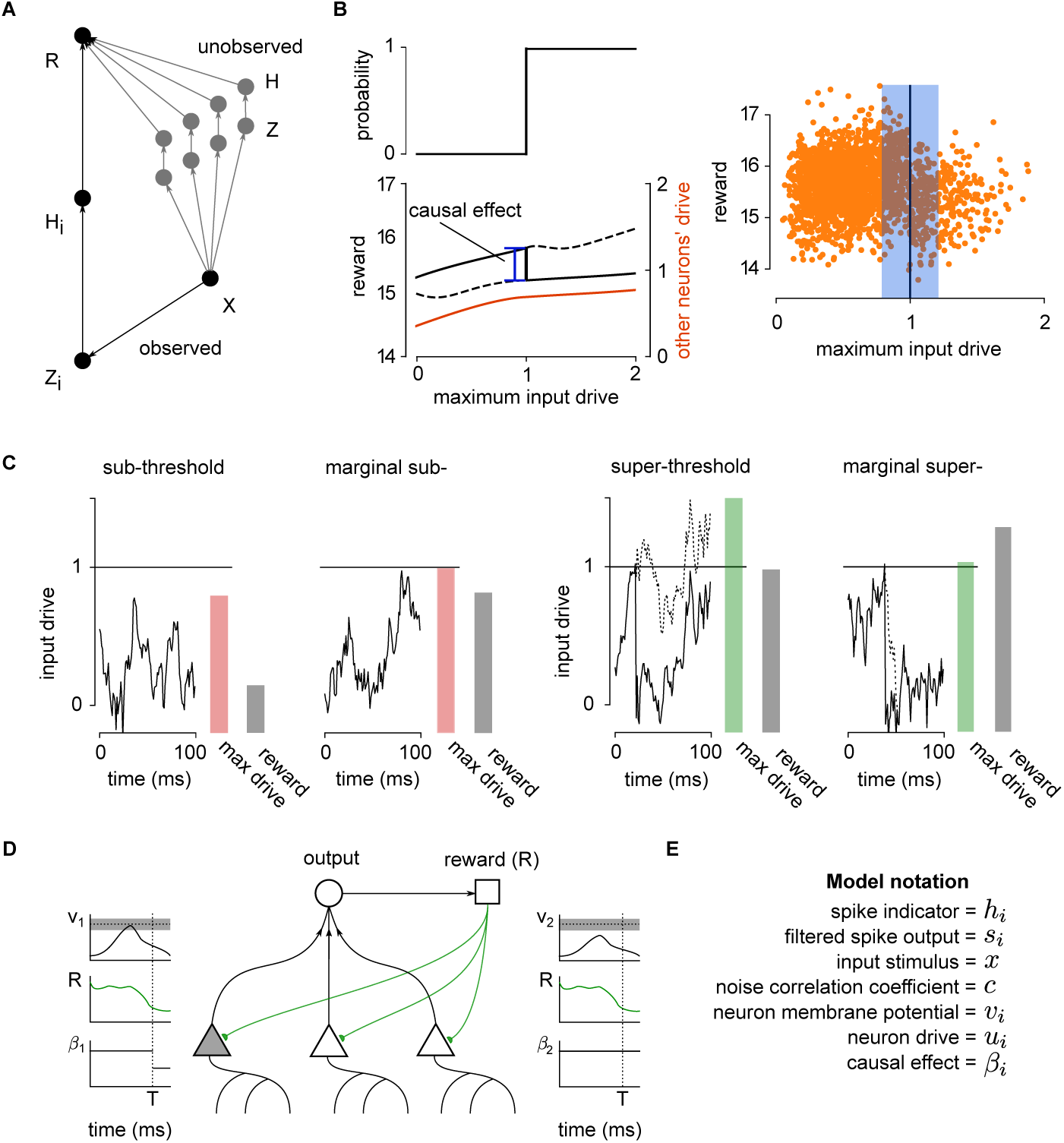
Using the spiking discontinuity for neural learning. (A) Graphical model describing neural network. Neuron *H*_*i*_ receives input *X*, which contributes to drive *Z*_*i*_. If drive is above the spiking threshold, then *H*_*i*_ is active. The activity contributes to reward *R*. Though not shown, this relationship may be mediated through downstream layers of a neural network, and complicated interactions with the environment. From neuron *H*_*i*_’s perspective, the activity of the other neurons *H* which also contribute to *R* is unobserved. (B) The reward may be tightly correlated with other neurons’ activity, which act as confounders. However any discontinuity in reward at the neuron’s spiking threshold can only be attributed to that neuron. The discontinuity at the threshold is thus a meaningful estimate of the causal effect (left). The effect of a spike on a reward function can be determined by considering data when the neuron is driven to be just above or just below threshold (right). (C) This is judged by looking at the neural drive to the neuron over a short time period. Marginal sub- and super-threshold cases can be distinguished by considering the maximum drive throughout this period. (D) Schematic showing how SDE operates in network of neurons. Each neuron contributes to output, and observes a resulting reward signal. Learning takes place at end of windows of length *T*. Only neurons whose input drive brought it close to, or just above, threshold (gray bar in voltage traces; compare neuron 1 to 2) update their estimate of *β*. (E) Model notation.

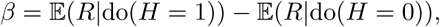

where do represents the do-operator, notation for an intervention. A causal effect can be measured through randomization, but sometimes can also be estimated without randomization by additionally observing the right variables to remove confounding. One such criterion is known as the back-door criterion [56]. In essence, outcomes are compared between treatment and non-treatment groups, when these additional variables satisfying the backdoor criterion are held fixed.

In a neural network setting, the causal effect of a neuron on reward can be seen, loosely, as the change in reward as a result of a change in the neuron’s activity made through randomization. Alternatively, it could be estimated by invoking the backdoor criterion and estimating the change in reward for a change in the neuron’s activity, when other neurons not downstream of that neuron are held fixed. In this spiking (binary) case the causal effect can be seen as a type of finite difference approximation of the partial derivative. That is, let *H*_*i*_ be the spiking indicator variable of a neuron *i* over a time window of length *T*, let *S*_*i*_ be a filtered version of the spiking activity, which contributes to determining the reward signal *R*. The neuron would like to estimate the effect of its spiking on *R*:

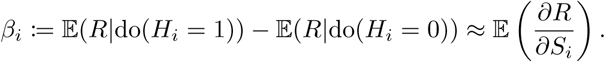

It can be shown that this quantity, under certain assumptions, does indeed approximate the reward gradient

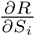 (see Methods). This establishes a link between causal inference and gradient-based learning, and suggests methods from causal inference may provide efficient algorithms to estimate reward gradients.

### Estimating a neuron’s causal effect using the spiking discontinuity

A discontinuity can be used to estimate a causal effect. We will refer it here as the Spiking Discontinuity Estimator (SDE). This is equivalent to the regression discontinuity design (RDD) approach which is popular in economics [34, 2]. For reasons outlined above, estimating the size of the discontinuity in an outcome at the threshold gives a measure of causal effect. As SDE derives from RDD it is valid when [34, 36]: first, whether a neuron spikes in an interval is not affected by the neuron’s spikes in that interval – there is no feedback over the interval; second, the spiking threshold is determined by intrinsic properties of the neuron, and not tuned for a specific drive; and third, noise in the network (and the fact that there are many presynaptic neurons) smooths out any discontinuities that may be a result of spiking in presynaptic inputs and thus, considering reward as a function of presynaptic drive, the neuron’s spiking is the only discontinuity observed. Under these assumptions neurons can produce unbiased estimates of causal effects using SDE.

For a neuron to apply SDE-based learning, it must track how close it is to spiking, whether it spiked or not, and observe the reward signal. More specifically, neurons spike when their maximal drive *Z*_*i*_ exceeds a threshold and then can receive feedback or a reward signal *R* through neuromodulator signals (Figure 1A). Then the comparison in reward between time periods when a neuron almost reaches its firing threshold to moments when it just reaches its threshold allows an SDE estimate of its own causal effect (Figure 1B,C,D,E).

To implement SDE, a neuron can estimate a piece-wise linear model of the reward function at time periods when its inputs place it close to threshold:

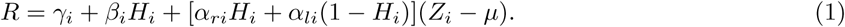

Here *H*_*i*_ is neuron *i*’s spiking indicator function, *γ*_*i*_, *α*_*li*_ and *α*_*ri*_ are the slopes that correct biases that would otherwise occur from having a finite bandwidth, *Z*_*i*_ is the maximum neural drive to the neuron over a short time period, and *β*_*i*_ represents the causal effect of neuron *i*’s spiking over a fixed time window of period *T*. The neural drive used here is the leaky, integrated input to the neuron, that obeys the same dynamics as the membrane potential except without a reset mechanism. By tracking the maximum drive attained over a short time period, marginally super-threshold inputs can be distinguished from well-above-threshold inputs, as required to apply SDE. Proposed physiological implementations of this model are described in the discussion.

To demonstrate that a neuron can use SDE to estimate causal effects here we analyze a simple two neuron network obeying leaky integrate-and-fire (LIF) dynamics. The neurons receive an input signal *x* with added noise, correlated with coefficient *c*. Each neuron weighs the noisy input by *w*_*i*_. The correlation in input noise induces a correlation in the output spike trains of the two neurons [66], thereby introducing confounding. The neural output determines a non-convex reward signal *R*. Most aspects of causal inference can be investigated in a simple model such as this [57], thus demonstrating that a neuron can estimate a causal effect with SDE in this simple case is an important first step to understanding how it can do so in a larger network.

Applying the SDE estimator shows that a neuron can estimate its causal effect (Figure 2A,B). To show how it removes confounding, we implement a simplified SDE estimator that considers only average difference in reward above and below threshold within a window of size *p*. When *p* is large this corresponds to the biased observed dependence estimator, while small *p* values approximate the SDE estimator and result in an unbiased estimate (Figure 2A). The window size *p* determines the variance of the estimator, as expected from theory [33]. Instead a locally linear SDE model, (1), can be used. This model is more robust to confounding (Figure 2B), allowing larger *p* values to be used. Thus the linear correction that is the basis of many RDD implementations [34] allows neurons to readily estimate their causal effect.

**Figure 2:**
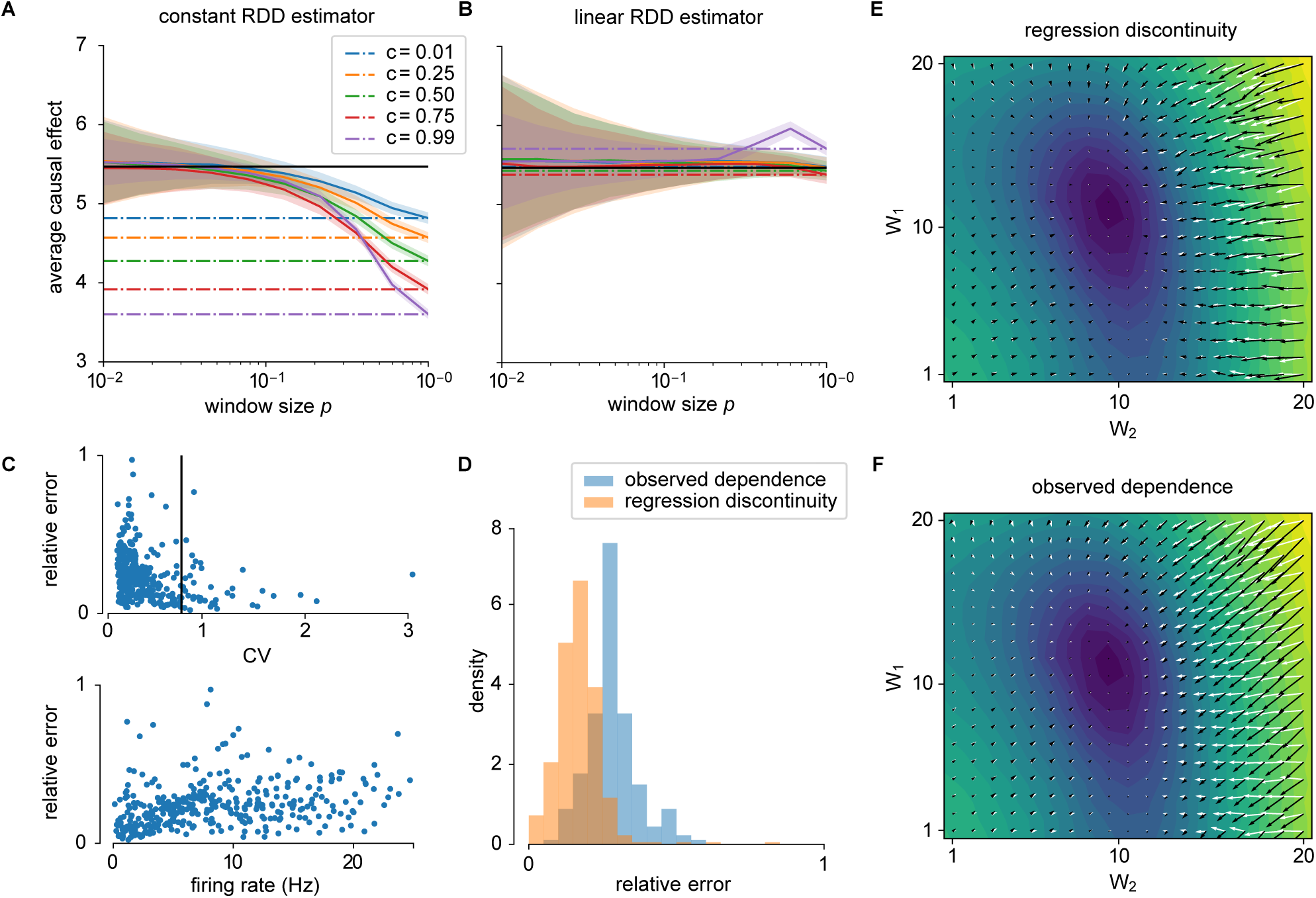
Estimating reward gradient with SDE in two-neuron network. (A) Estimates of causal effect (black line) using a constant SDE model (difference in mean reward when neuron is within a window *p* of threshold) reveals confounding for high *p* values and highly correlated activity. *p* = 1 represents the observed dependence, revealing the extent of confounding (dashed lines). Curves show mean plus/minus standard deviation over 50 simulations. (B) The linear SDE model is unbiased over larger window sizes and more highly correlated activity (high *c*). (C) Relative error in estimates of causal effect over a range of weights (1 ≤ *w*_*i*_ ≤ 20) show lower error with higher coefficient of variability (CV; top panel), and lower error with lower firing rate (bottom panel). (D) Over this range of weights, SDE estimates are less biased than just the naive observed dependence. (E,F) Approximation to the reward gradient overlaid on the expected reward landscape. The white vector field corresponds to the true gradient field, the black field correspond to the SDE (E) and OD (F) estimates. The observed dependence is biased by correlations between neuron 1 and 2 – changes in reward caused by neuron 1 are also attributed to neuron 2.

To investigate the robustness of the SDE estimator, we systematically vary the weights, *w*_*i*_, of the network. SDE works better when activity is fluctuation-driven and at a lower firing rate (Figure 2C). Thus SDE is most applicable in irregular but synchronous activity regimes [12]. Over this range of network weights SDE is less biased than the observed dependence (Figure 2D).

### An discontinuity-based learning rule

A canonical causal inference problem is learning – in order to update synaptic weights to improve performance a neuron needs to know its causal effect on that performance. The spiking discontinuity allows neurons to estimate their causal effect. We can show that this provides a rule for neurons to update their weights to maximize a reward. For gradient-based learning, to do this the neuron must relate the gradient 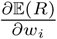 to something involving the causal effect. One complication from a theoretical standpoint is that the distribution over which the above expectation is taken depends on the parameters *w*, meaning the derivative cannot be immediately transferred inside the expectation. Additional assumptions are needed to justify something like this. It can be shown (see Methods) that under the following assumptions:

1. The neural network parameters only affect the expected reward through their spiking activity, meaning that 𝔼(*R*|*H*) is independent of parameters **w**.
2. The gradient term 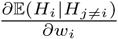 is independent of *H*_*j*≠*i*_.
3. Neurons *H*_*j*≠*i*_ satisfy the backdoor criterion with respect to *H*_*i*_ → *R*.

then:

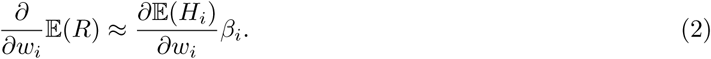

These assumptions are not necessarily true for the dynamical networks considered here, and their validity must be tested empirically. They were shown to be reasonable over certain activity regimes in the simulations used here (Supplementary Material). Thus the estimate of causal effect can be used to update synaptic weights using a gradient-descent based rule.

To demonstrate how a neuron can learn *β* through SDE, we derive an online learning rule from the locally linear SDE model. The rule takes the form:

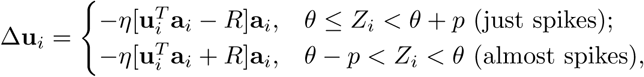

where **u**_*i*_ are the parameters of the linear model required to estimate *β*_*i*_, *η* is a learning rate, and **a**_*i*_ are drive-dependent terms (see Methods). This rule can then be applied along with Equation (2) to update synaptic weights. The causal effect can be used to estimate 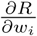 (Figure 2E,F), and thus the SDE estimator may be used in a learning rule to update weights so as to maximize expected reward.

When applied to the toy network, the online learning rule (Figure 3A) estimates *β* over the course of seconds (Figure 3B). When the estimated *β* is then used to update weights to maximize expected reward in an unconfounded network (uncorrelated – *c* = 0.01), SDE-based learning exhibits higher variance than learning using the observed dependence. SDE-based learning exhibits trajectories that are initially meander while the estimate of *β* settles down (Figure 3C). When a confounded network (correlated – *c* = 0.5) is used SDE exhibits similar performance, while learning based on the observed dependence sometimes fails to converge due to the bias in gradient estimate. In this case SDE-based learning also converges faster than learning based on observed dependence (Figure 3D,E). Thus the SDE-based learning rule allows a network to be trained on the basis of confounded inputs.

**Figure 3:**
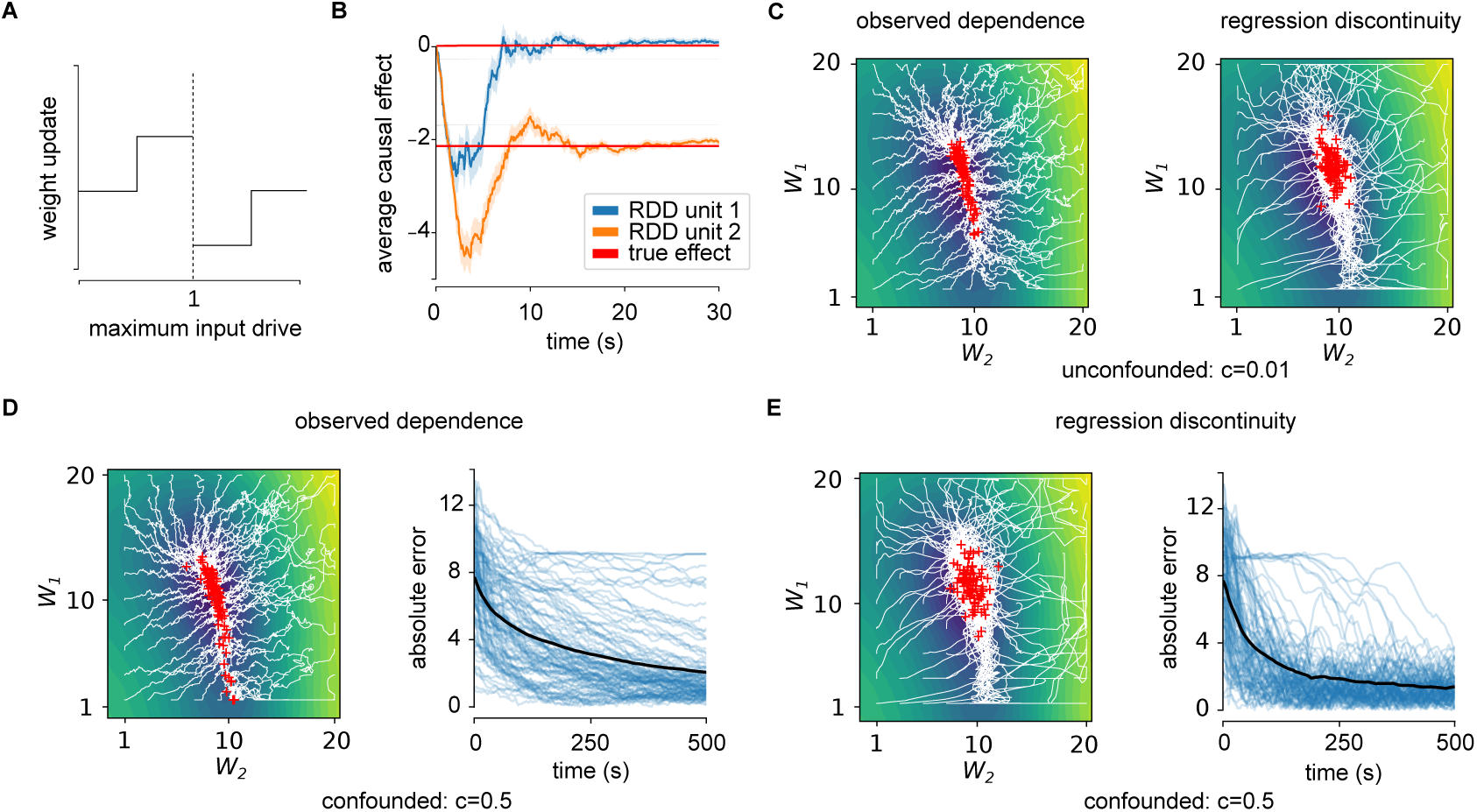
Applying the discontinuity learning rule. (A) Sign of SDE learning rule updates are based on whether neuron is driven marginally below or above threshold. (B) Applying rule to estimate *β* for two sample neurons shows convergence within 10s (red curves). Error bars represent standard error of the mean over 50 simulations. (C) Convergence of observed dependence (left) and SDE (right) learning rule to unconfounded network (*c* = 0.01). Observed dependence converges more directly to bottom of valley, while SDE trajectories have higher variance. (D,E) Convergence of observed dependence (D) and SDE (E) learning rule to confounded network (*c* = 0.5). Right panels: error as a function of time for individual traces (blue curves) and mean (black curve). With confounding learning based on observed dependence converges slowly or not at all, whereas SDE succeeds.

### Application to BCI learning

To demonstrate the behavior of SDE learning in a more realistic setting, we consider learning with intracortical brain-computer interfaces (BCIs) [19, 61, 44, 26, 51]. BCIs provide an excellent opportunity to test theories of learning and to understand how neurons solve causal inference problems. This is because in a BCI the exact mapping from neural activity to behavior and reward is known by construction [26] – the true causal effect of each neuron is known. Recording from neurons that directly determine BCI output as well as those that do not allows us to observe if and how neural populations distinguish a causal relationship to BCI output from a correlational one. Here we compare SDE-based learning and observed dependence-based learning to known behavioral results in a setting that involves correlations among neurons. We focus on single-unit BCIs [51], which map the activity of individual units to cursor motion. This is a meaningful test for SDE-based learning because the small numbers of units involved in the output function mean that a single spike has a measurable effect on the output.

We focus in particular on a type of BCI called a dual-control BCI, which requires a user control a BCI and simultaneously engage in motor control [6, 54, 50, 39] (Figure 4A). Dual-control BCIs engage primary motor units in a unique way: the units directly responsible for the cursor movement (henceforth control units) change their tuning and effective connectivity properties to control an output, while other observed units retain their association to motor control (Figure 4B) [50, 39]. In other words, control units modify their activity to perform a dual-control BCI task while other observed neurons, whose activity is correlated with the control neurons, do not. The issue of confounding is thus particularly relevant in dual-control BCIs.

We run simulations inspired by learning in a dual-control BCI task. Ten LIF neurons are correlated with one another, representing a population of neurons similarly tuned to wrist motion. A cost function is defined that requires the control unit to reach a target firing rate, related to the BCI output, and the remainder of the units to reach a separate firing rate, related to their role motor control. We observe that, using the SDE-based learning rule, the neurons learn to minimize this cost function – the control unit changes its weights specifically. This performance is independent of the degree of correlation between the neurons (Figure 4C). An observed-dependence based learning rule also achieves this, but the final performance depends on the degree of correlation (Figure 4D). Yet dual-control BCI studies have shown that performance is independent of a control unit’s tuning to wrist motion or effective connectivity with other units [50, 39]. Examination of the synaptic weights as learning proceeds shows the control unit in SDE-based learning quickly separates itself from the other units (Figure 4E). The control unit’s weight in observed-dependence based learning, on the other hand, initially increases with the other units (Figure 4F). It cannot distinguish its effect on the cost function from the other units’ effect, even with relatively low, physiologically relevant correlations (e.g. *c* = 0.3, [15]). SDE-based learning may be relevant in the context of BCI learning.

**Figure 4:**
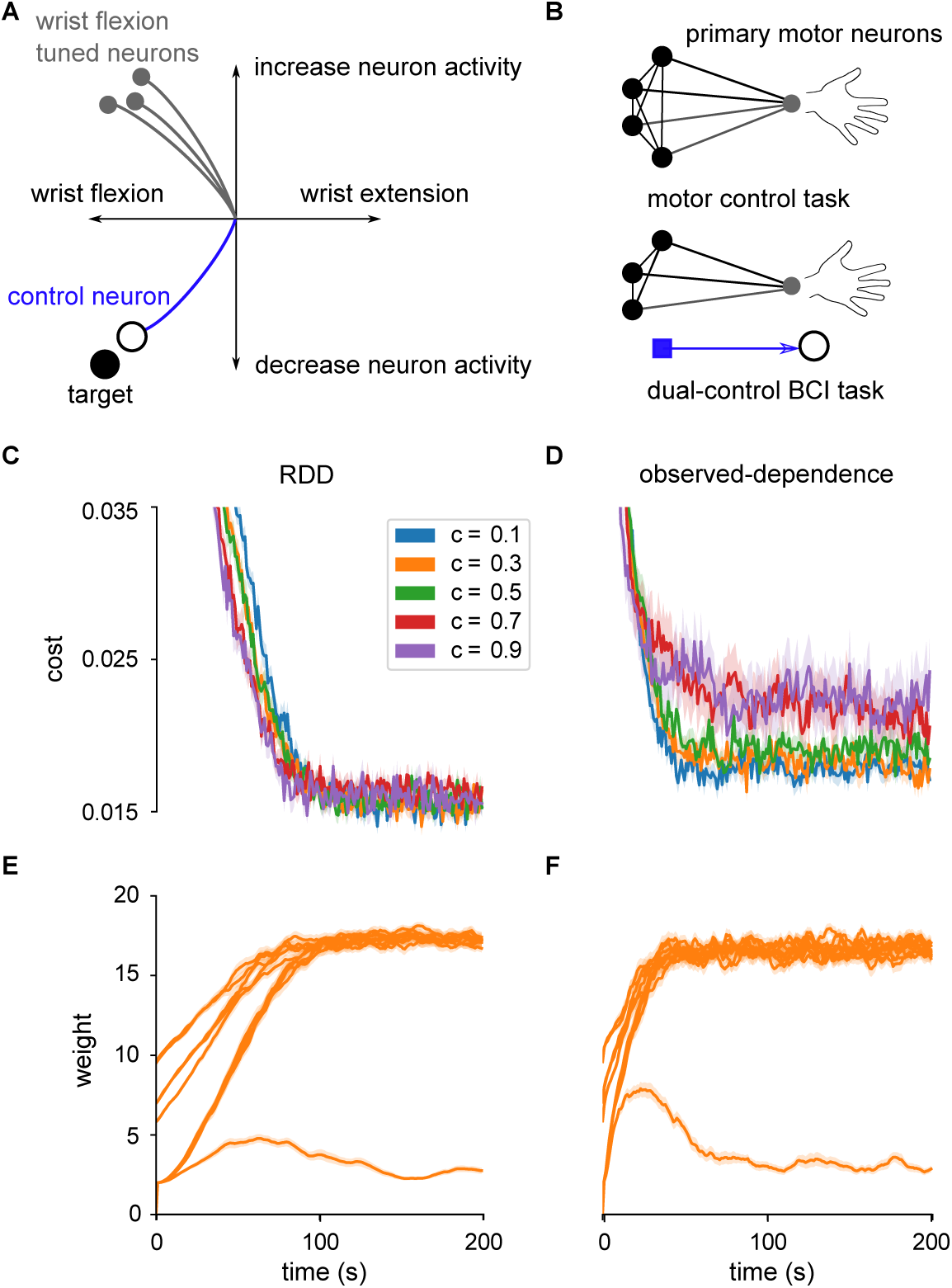
Application to BCI learning. (A) Dual-control BCI setup [50]. Vertical axis of a cursor is controlled by one control neuron, horizontal axis is controlled by wrist flexion/extension. Wrist flexion-tuned neurons (gray) would move cursor up and right when wrist is flexed. To reach a target, task may require the control neuron (blue) dissociate its relation to wrist motion – decrease its activity and flex wrist simultaneously – to move cursor (white cursor) to target (black circle). (B) A population of primary motor neurons correlated with (tuned to) wrist motion. One neuron is chosen to control the BCI output (blue square), requiring it to fire independently of other neurons, which maintain their relation to each other and wrist motion [39]. (C) SDE-based learning of dual-control BCI task for different levels of correlation. Curves show mean over 10 simulations, shaded regions indicate standard error. (D) Observed dependence-based learning of dual-control BCI task. (E) Synaptic weights for each unit with *c* = 0.3 in SDE-based learning. Control unit is single, smaller weight. (F) Synaptic weights for each unit in observed-dependence based learning.

## Discussion

Here we have cast neural learning explicitly as a causal inference problem, and have shown that neurons can estimate their causal effect using their spiking mechanism. In this way we found that spiking can be an advantage, allowing neurons to quantify their causal effects in an unbiased way. We outlined sufficient assumptions for this causal inference to be correct, and have shown how this causal inference estimate can be turned into a learning rule that can optimize the synaptic weights of a neuron.

The rule is inspired by the regression discontinuity design commonly used in econometrics [3]. Previous work in econometrics has considered how the threshold may be changed [17] and how that can be used to optimize utility [47]. We have extended this work by deriving a rule that allows for the neuron to change its weights, instead of just its threshold, to increase utility. This provides the ability to change not just the threshold but the construction of the variable being thresholded to optimize reward.

It is important to note that other neural learning rules also perform causal inference. Thus the SDE-based learning rule can be placed in the context of other neural learning mechanisms. First, as many authors have noted, any reinforcement learning algorithm relies on estimating the effect of an agent’s/neuron’s activity on a reward signal. Learning by operant conditioning relies on learning a causal relationship (compared to classical conditioning, which only relies on learning a correlation) [76, 56, 28, 25]. Causal inference is, at least implicitly, the basis of reinforcement learning.

There is a large literature on how reinforcement learning algorithms can be implemented in the brain. It it well known there are many neuromodulators which may represent reward or expected reward, including dopaminergic neurons from the substantia nigra to the ventral striatum representing a reward prediction error [75, 62]. Many of these methods use something like the REINFORCE algorithm [74], a policy gradient method in which locally added noise is correlated with reward and this correlation is used to update weights. This gives an unbiased estimate of the causal effect because the noise is assumed to be independent, private to each neuron. These ideas have extensively been used to model learning in brains [11, 21, 20, 44, 49, 77, 64]. Learning in birdsong is a particularly well developed example of this form of learning [20]. In birdsong learning in zebra finches, neurons from area LMAN synapse onto neurons in area RA. These synapses are referred to as ‘empiric’ synapses, and are treated by the neurons as an ‘experimenter’, producing random perturbations which can be used to estimate causal effects. This is a compelling account of learning in birdsong, however it relies on the specific structural form of the learning circuit. It is unknown more generally how a neuron may estimate what is perturbative noise without these structural specifics, and thus if it can provide an account of learning more generally.

There are two factors that cast doubt on the use of reinforcement learning-type algorithms broadly in neural circuits. First, even for a fixed stimulus, noise is correlated across neurons [5, 15, 37, 7, 78]. Thus if the noise a neuron uses for learning is correlated with other neurons then it can not know which neuron’s changes in output is responsible for changes in reward. In such a case, the synchronizing presynaptic activity acts as a confounder. Thus, as discussed, such algorithms require biophysical mechanisms to distinguish independent perturbative noise from correlated input signals in presynaptic activity, and in general it is unclear how a neuron could can do this. And, second, learning with perturbations scales poorly with network size [64, 20, 60]. Thus neurons may use alternatives to these reinforcement-learning algorithms.

Given the inefficiency of these reinforcement learning algorithms, a number of authors have looked to learning in artificial neural networks for inspiration. In artificial neural networks, the credit assignment problem is efficiently solved using the backpropagation algorithm, which allows efficiently calculating gradients. Backpropagation requires differentiable systems, which spiking neurons are not. Indeed, cortical networks often have low firing rates in which the stochastic and discontinuous nature of spiking output cannot be neglected [65]. It also requires full knowledge of the system, which is often not the case if parts of the system relate to the outside world. No known structures exist in the brain that could exactly implement backpropagation. Yet, backpropagation is significantly more powerful than perturbation-based methods – it is the only known algorithm able to solve large-scale problems at a human-level [42]. The success of backpropagation suggests that efficient methods for computing gradients are needed for solving large-scale learning problems.

A number of recent approaches have looked for neural mechanisms which allow for more specific feedback signals to support learning, inspired by the efficiency of the backpropagation algorithm [27, 38, 45]. By providing neuron-specific error feedback, these methods provide information about each neuron’s causal effect. In particular, this work has proposed neural learning in cortex uses neuron-specific feedback by separating the computation into two compartments. In one component the neuron integrates feedforward inputs, and in the other component the neuron integrates feedback signals in order to estimate something like a reward gradient. This separation is plausibly implementable in apical and basal dendritic compartments of cortical pyramidal neurons. This approach is shared with SDE-based learning, in which the casual effect is estimated separately and then used to drive learning. We may thus expect that an SDE-based learning rule could be readily combined with these compartmental models of learning in cortex to provide a biologically plausible account of efficient learning.

Finally, it must be noted how the learning rule derived here relates to the dominant spike-based learning paradigm – spike timing dependent plasticity (STDP [10]). STDP performs unsupervised learning, so is not directly related to the type of optimization considered here. Reward-modulated STDP (R-STDP) can be shown to approximate the reinforcement learning policy gradient type algorithms described above [46, 24, 35]. Thus R-STDP can be cast as performing a type of causal inference on a reward signal, and shares the same features and caveats as outlined above. Thus we see that learning rules that aim at maximizing some reward either implicitly or explicitly involve a neuron estimating its causal effect on that reward signal. Explicitly recognizing this can lead to new methods and understanding.

### Caveats and advantages

There are multiple caveats for the use of an SDE-based learning rule. The first is that the rule is only applicable in cases where the effect of a single spike is relevant. Depending on the way a network is constructed, the importance of each neuron may decrease as the size of the network is increased. As the influence of a neuron vanishes, it becomes hard to estimate this influence. While this general scaling behavior is shared with other algorithms (e.g. backpropagation), it is more crucial for SDE where there will be some noise in the evaluation of the outcome.

A second caveat is that the SDE rule does not solve the temporal credit assignment problem. It requires us to know which output is associated with which kind of activity. There are multiple approaches that can help solve this kind of problem, including actor-critic methods and eligibility traces from reinforcement learning theory [68].

A third caveat is that, as implemented here, the rule learns the effect of a neuron’s activity on a reward signal for a fixed input. Thus the rule is applicable in cases where a fixed output is required. This includes learning stereotyped motor actions, or learning to mimic a parent’s birdsong [20, 21]. Applying the learning rule to networks with varying inputs, as in many supervised learning tasks, would require extensions. One possible extension that may address these caveats is, rather than directly estimate the effect of a neuron’s output on a reward function, to use the method to learn weights on feedback signals so as to approximate the causal effect – that is, to use the SDE rule to “learn how to learn”[40]. This approach has been shown to work in artificial neural networks, suggesting SDE-based learning may provide a biologically plausible and scale-able approach to learning [16, 48, 27].

This paper introduces the SDE method to neuronal learning and artificial neural networks. It illustrates the difference in behavior of SDE and observed-dependence learning in the presence of confounding. While it is of a speculative nature, at least in cases where reward signals are observed, it does provide a biologically plausible account of neural learning. It addresses a number of issues with other learning mechanisms.

SDE-based learning does not require independent noise. It is sufficient that something, in fact anything that is presynaptic, produce variability. As such, SDE approaches do not require the noise source to be directly measured. This allows to rule to be applied in a wider range of neural circuits or artificial systems.

SDE-based learning removes confounding due to noise correlations. Noise correlations are known to be significant in many sensory processing areas [15]. While noise correlations’ role in sensory encoding has been well studied [5, 15, 37, 7, 78], their role in learning has been less studied. This work suggests that understanding learning as a causal inference problem can provide insight into the role of noise correlations in learning.

Finally, in a lot of theoretical work, spiking is seen as a disadvantage, and models thus aim to remove spiking discontinuities through smoothing responses [32, 31, 43]. The SDE rule, on the other hand, exploits the spiking discontinuity. Moreover, the rule can operate in environments with non-differentiable or discontinuous reward functions. In many real-world cases, gradient descent would be useless: even if the brain could implement it, the outside world does not provide gradients (but see [73]). Our approach may thus be useful even in scenarios, such as reinforcement learning, where spiking is not necessary. Spiking may, in this sense, allow a natural way of understanding a neuron’s causal influence in a complex world.

### Compatibility with known physiology

If neurons perform something like SDE-based learning we should expect that they exhibit certain physiological properties. We thus want to discuss the concrete demands of SDE-based learning and how they relate to past experiments.

The SDE learning rule is applied only when a neuron’s membrane potential is close to threshold, regardless of spiking. This means inputs that place a neuron close to threshold, but do not elicit a spike, still result in plasticity. This type of sub-threshold dependent plasticity is known to occur [22, 23, 67]. This also means that plasticity will not occur for inputs that place a neuron too far below threshold. In past models of voltage-dependent plasticity and experimental findings, changes do not occur when postsynaptic voltages are too low (Figure 5A) [14, 4]. And the SDE learning rule predicts that plasticity does not occur when postsynaptic voltages are too high. However, in many voltage-dependent plasticity models, potentiation does occur for inputs well-above the spiking threshold. But, as SDE-based learning occurs in low-firing rate regimes, inputs that place a neuron too far above threshold are rare. So this discrepancy may not be relevant. Thus threshold-adjacent plasticity as required for SDE-based learning appears to be compatible with neuronal physiology.

**Figure 5:**
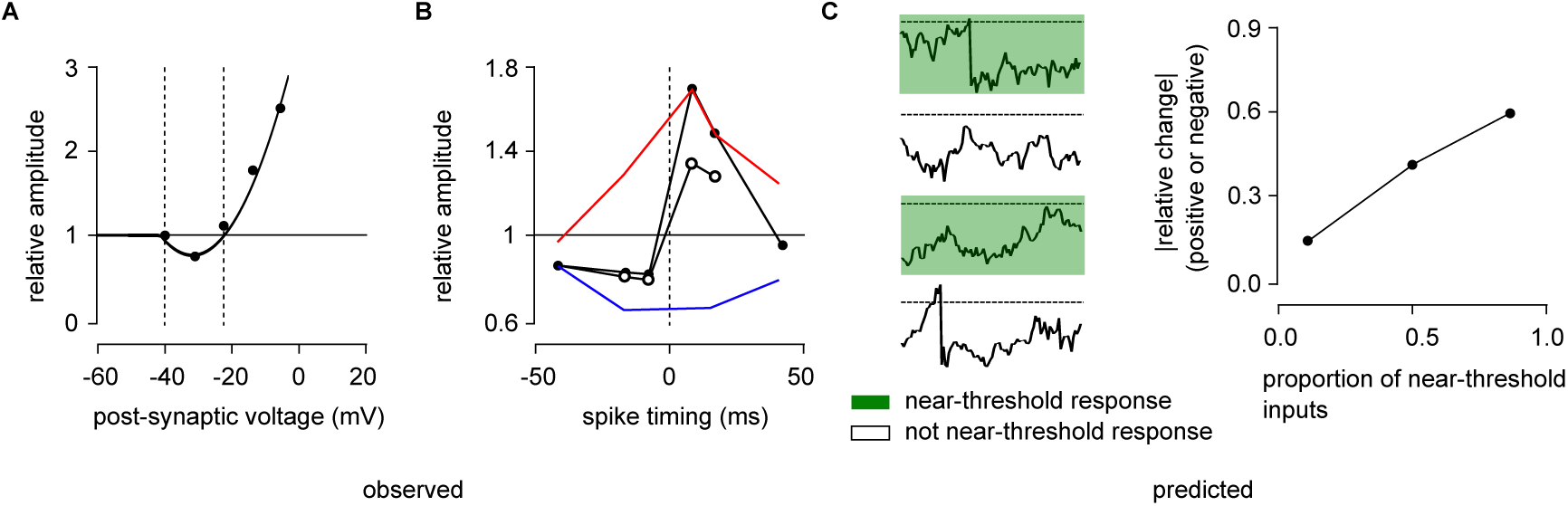
Feasibility and predictions of SDE-based learning. (A) Voltage-clamp experiments in adult mice CA1 hippocampal neurons [53]. Amplitude of potentiation or depression depends on post-synaptic voltage. (B) Neuromodulators affect magnitude of STDP. High proportion of *β*-adrenergic agonist to M1 muscarinic agonist (filled circles), compared with lower proportion (unfilled circles) in visual cortex layer II/III neurons. Only *β*-adrenergic agonist produces potentiation (red curve), and only M1 muscarinic agonist produces depression (blue) [63]. (C) Prediction of SDE-based learning. Over a fixed time window a reward is administered when neuron spikes. Stimuli are identified which place the neuron’s input drive close to spiking threshold. SDE-based learning predicts an increase synaptic changes for a set of stimuli containing a high proportion of near threshold inputs, but that keeps overall firing rate constant.

The SDE-based learning predicts that spiking switches the sign of plasticity. Some experiments and phenomenological models of voltage-dependent synaptic plasticity do capture this behavior [4, 14, 13]. In these models, plasticity changes from depression to potentiation near the spiking threshold (Figure 5A). Thus this property of SDE-based learning also appears to agree with some models and experimental findings.

The SDE-based learning is dependent on neuromodulation. Neuromodulated-STDP is well studied in models [24, 63, 55]. However, SDE-based learning requires plasticity switch sign under different levels of reward. This may be communicated by neuromodulation. There is some evidence that the relative balance between adrenergic and M1 muscarinic agonists alters both the sign and magnitude of STDP in layer II/III visual cortical neurons [63] (Figure 5B). To the best of our knowledge, how such behavior interacts with postsynaptic voltage dependence as required by SDE-learning is unknown. Thus, taken together, these factors show SDE-based learning may well be compatible with known neuronal physiology, and leads to predictions that can demonstrate SDE-based learning.

### Experimental predictions

A number of experiments can be used to test if neurons use SDE to perform causal inference. This has two aspects, neither of which have been explicitly tested before: (1) do neurons learn in the presence of strong confounding, which is do causal inference and (2) do they use SDE to do so? To test both of these we propose scenarios that let an experimenter control the degree of confounding in a population of neurons’ inputs and control a reward signal.

To test if neurons learn in the presence of strong confounding, we propose an experiment where a lot of neurons are stimulated in a time-varying way but only one of the neurons has a causal influence, which is defined in a brain computer interface (BCI) experiment (like [18]). Due to the joint stimulation, all neurons will be correlated at first and therefore all neurons will correlate with the reward. But as only one neuron has a causal influence, only that neuron should increase its firing (as in Fig. 4. Testing how each neuron adjusts its firing properties in the presence of different strengths of correlation would establish if the neuron causally related to the reward signal can be distinguished from the others.

A similar experiment can also test if neurons specifically use SDE to achieve deconfounding. SDE-based learning happens when a neuron is close to threshold. We could use manipulations to affect when a neuron is close to threshold, e.g. through patch clamping or optical stimulation. By identifying these inputs, and then varying the proportion of trials in which a neuron is close to threshold, while keeping the total firing rate the same, the SDE hypothesis can be tested (Figure 5C). SDE-based learning predicts that a set of trials that contain many near-threshold events will result in faster learning than in a set of trials that place a neuron either well above or well below threshold.

Alternatively, rather than changing input activity to bring a neuron close to threshold, or not, SDE-based learning can be tested by varying when reward is administered. Again, identifying trials when a neuron is close to threshold and introducing a neurotransmitter signalling reward (e.g. dopamine) at the end of these trials should, according to SDE learning, result in an increased learning rate, compared to introducing the same amount of neurotransmitter whenever the neuron spiked. These experiments are important future work that would demonstrate SDE-based learning.

### Conclusion

The most important aspect of SDE-based learning is the explicit focus on causality. A causal model is one that can describe the effects of an agent’s actions on an environment. Learning through the reinforcement of an agent’s actions relies, even if implicitly, on a causal understanding of the environment [25, 41]. Here, by explicitly casting learning as a problem of causal inference we have developed a novel learning rule for spiking neural networks. We present the first model to propose a neuron does causal inference. We believe that focusing on causality is essential when thinking about the brain or, in fact, any system that interacts with the real world.

## Methods

### Neuron, noise and reward model

We consider the activity of a network of *n* neurons whose activity is described by their spike times

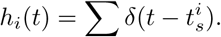

Here *n* = 2. Synaptic dynamics **s** *∈* ℝ^*n*^ are given by

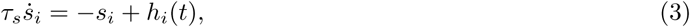

for synaptic time scale *τ*_*s*_. An instantaneous reward is given by *R*(**s**) ∈ ℝ. In order to have a more smooth reward signal, *R* is a function of **s** rather than **h**. The reward function used here has the form of a Rosenbrock function:

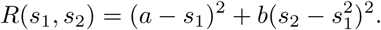

The neurons obey leaky integrate-and-fire (LIF) dynamics

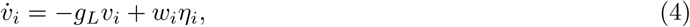

where integrate and fire means simply:

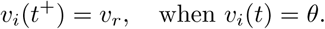

Noisy input *η*_*i*_ is comprised of a common DC current, *x*, and noise term, *ξ*(*t*), plus an individual noise term, *ξ*_*i*_(*t*):

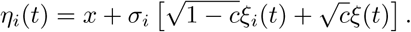

The noise processes are independent white noise: 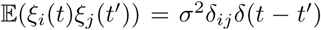. This parameterization is chosen so that the inputs *η*_1,2_ have correlation coefficient *c*. Simulations are performed with a step size of Δ*t* = 1ms. Here the reset potential was set to *v*_*r*_ = 0. Borrowing notation from Xie and Seung 2004 [77], the firing rate of a noisy integrate and fire neuron is

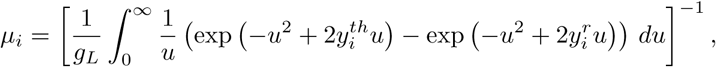

where 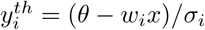 and 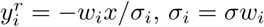 is the input noise standard deviation.

We define the input drive to the neuron as the leaky integrated input without a reset mechanism. That is, over each simulated window of length *T* :

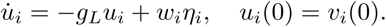

The SDE method operates when a neuron receives inputs that place it close to its spiking threshold – either nearly spiking or barely spiking – over a given time window. In order to identify these time periods, the method uses the maximum input drive to the neuron:

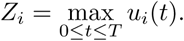

The input drive is used here instead of membrane potential directly because it can distinguish between marginally super-threshold inputs and easily super-threshold inputs, whereas this information is lost in the voltage dynamics once a reset occurs. Here a time period of *T* = 50ms was used. Reward is administered at the end of this period: *R* = *R*(**s**_*T*_).

### Policy gradient methods in neural networks

The dynamics given by (4) generate an ergodic Markov process with a stationary distribution denoted *ρ*. We consider the problem of finding network parameters that maximize the expected reward with respect to *ρ*. In reinforcement learning, performing optimization directly on the expected reward leads to policy gradient methods [69]. These typically rely on either finite difference approximations or a likelihood-ratio decomposition [74]. Both approaches ultimately can be seen as performing stochastic gradient descent, updating parameters by approximating the expected reward gradient:

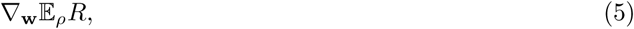

for neural network parameters **w**.

Here capital letters are used to denote the random variables drawn from the stationary distribution, corresponding to their dynamic lower-case equivalent above. Density *ρ* represents a joint distribution over variables (*Z, H, S, R*), the maximum input drive, spiking indicator function, filtered spiking output, and reward variable, respectively. The spiking indicator function is defined as *H*_*i*_ = 𝕀 (*Z*_*i*_ ≥ *θ*), for threshold *θ*.

We wish to evaluate (5). In general there is no reason to expect that taking a derivative of an expectation with respect to some parameters will have the same form as the corresponding derivative of a deterministic function. However in some cases this is true, for example when the parameters are separable from the distribution over which the expectation is taken (sometimes relying on what is called the reparameterization trick [60, 29]). Here we show that, even when the reparameterization trick is unavailable, if the system contains a Bernoulli (spiking indicator) variable then the expression for the reward gradient also matches a form we might expect from taking the gradient of a deterministic function.

The expected reward can be expressed as

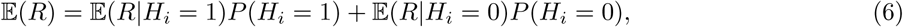

for a neuron *i*. The key assumption we make is the following:

#### Assumption 1.

The neural network parameters only affect the expected reward through their spiking activity, meaning that 𝔼(*R*|*H*) is independent of parameters **w**.

We would like to take derivatives of 𝔼(*R*) with respect to **w** using (6). However even if, under Assumption 1, the joint conditional expectation 𝔼(*R*|*H*) is independent of **w**, it is not necessarily the case that the marginal conditional expectation, 𝔼(*R*|*H*_*i*_), is independent of **w**. This is because it involves marginalization over unobserved neurons *H*_*j*≠*i*_, which may have some relation to *H*_*i*_ that is dependent on **w**. That is,

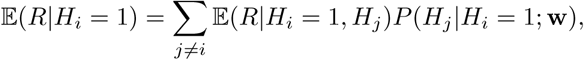

where the conditional probabilities may depend on **w**. This dependence complicates taking derivatives of the decomposition (6) with respect to **w**.

#### Unconfounded network

We can gain intuition about how to proceed by first making a simplifying assumption. If we assume that *H*_*i*_ ***╨*** *H*_*j*≠*i*_, then the reward gradient is simple to compute from (6). This is because now the parameter *w*_*i*_ only affects the probability of neuron *i* spiking:

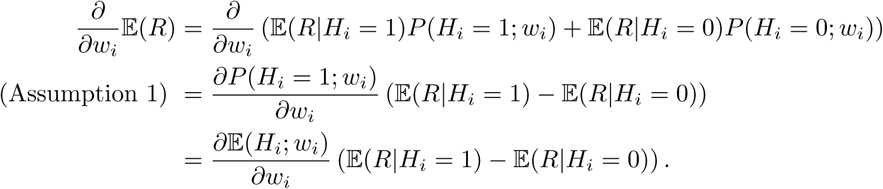

This resembles a type of finite difference estimate of the gradient we might use if the system were deterministic and *H* were differentiable:

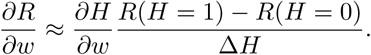

Based on the independence assumption we call this the unconfounded case. In fact the same decomposition is utilized in a REINFORCE-based method derived by Seung 2003 [64].

#### Confounded network

Generally, of course, it is not the case that *H*_*i*_ ***╨*** *H*_*j*≠*i*_, and then we must decompose the expected reward into:

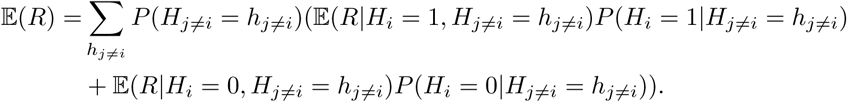

This means the expected reward gradient is given by:

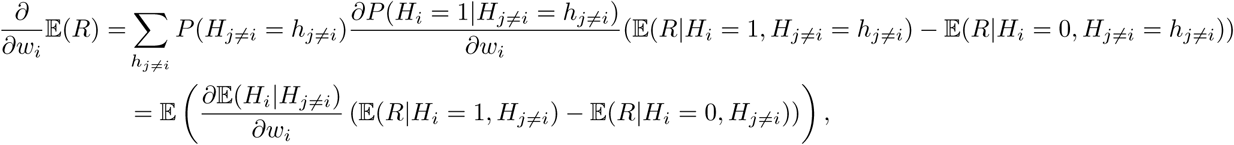

again making use of Assumption 1. We additionally make the following approximation:

##### Assumption 2.

The gradient term 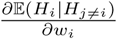 is independent of *H*_*j*≠*i*_.

This means we can move the gradient term out of the expectation to give:

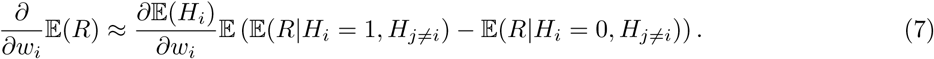

We assume that how the neuron’s activity responds to changes in synaptic weights, 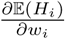, is known by the neuron. Thus it remains to estimate 𝔼(*R*|*H*_*i*_ = 1, *H*_*j*≠*i*_) − 𝔼(*R*|*H*_*i*_ = 0, *H*_*j*≠*i*_). It would seem this relies on a neuron observing other neurons’ activity. Below we show how it can be estimated, however, using methods from causal inference.

### The unbiased gradient estimator as a causal effect

We can identify the unbiased estimator (Equation (7)) as a causal effect estimator. To understand precisely what this means, here we describe a causal model.

A causal model is a Bayesian network along with a mechanism to determine how the network will respond to intervention. This means a causal model is a directed acyclic graph (DAG) *𝒢* over a set of random variables 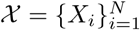 and a probability distribution *P* that factorizes over *𝒢* [56].

An intervention on a single variable is denoted do(*X*_*i*_ = *y*). Intervening on a variable removes the edges to that variable from its parents, Pa_*X*_*i*, and forces the variable to take on a specific value: 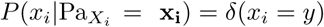. Given the ability to intervene, the average treatment effect (ATE), or causal effect, between an outcome variable *X*_*j*_ and a binary variable *X*_*i*_ can be defined as:

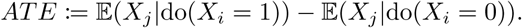

We make use of the following result: if **S**_*ij*_ *⊂ 𝒳* is a set of variables that satisfy the *back-door criteria* with respect to *X*_*i*_ → *X*_*j*_, then it satisfies the following: (i) **S**_*ij*_ blocks all paths from *X*_*i*_ to *X*_*j*_ that go into *S*_*i*_, and (ii) no variable in **S**_*ij*_ is a descendant of *X*_*i*_. In this case the interventional expectation can be inferred from

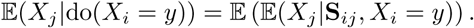

Given this framework, here we will define the causal effect of a neuron as the average causal effect of a neuron *H*_*i*_ spiking or not spiking on a reward signal, *R*:

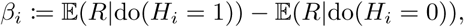

where *H*_*i*_ and *R* are evaluated over a short time window of length *T*.

We make the final assumption:

#### Assumption 3.

Neurons *H*_*j*≠*i*_ satisfy the backdoor criterion with respect to *H*_*i*_ → *R*.

Then it is the case that the reward gradient estimator, (7), in fact corresponds to:

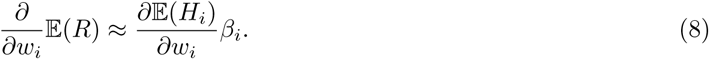

Thus we have the result that, in a confounded, spiking network, gradient descent learning corresponds to causal learning. The validity of the three above assumptions for a network of integrate and fire neurons is demonstrated in the Supplementary Material (Section S1). The relation between the causal effect and a finite difference estimate of the reward gradient is presented in the Supplementary Material (Section S2).

### Using the spiking discontinuity

To remove confounding, SDE considers only the marginal super- and sub-threshold periods of time to estimate (8). This works because the discontinuity in the neuron’s response induces a detectable difference in outcome for only a negligible difference between sampled populations (sub- and super-threshold periods). The SDE method estimates [34]:

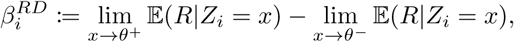

for maximum input drive obtained over a short time window, *Z*_*i*_, and spiking threshold, *θ*; thus, *Z*_*i*_ *< θ* means neuron *i* does not spike and *Z*_*i*_ *≥θ* means it does.

To estimate 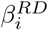, a neuron can estimate a piece-wise linear model of the reward function:

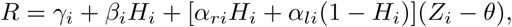

locally, when *Z*_*i*_ is within a small window *p* of threshold. Here *γ*_*i*_, *α*_*li*_ and *α*_*ri*_ are nuisance parameters, and *β*_*i*_ is the causal effect of interest. This means we can estimate 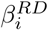 from

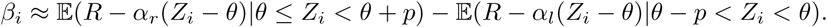

A neuron can learn an estimate of 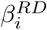 through a least squares minimization on the model parameters *β*_*i*_, *α*_*l*_, *α*_*r*_. That is, if we let **u**_*i*_ = [*β*_*i*_, *α*_*r*_, *α*_*l*_]^*T*^ and **a**_*t*_ = [1, *h*_*i,t*_(*z*_*i,t*_ *− θ*), (1 − *h*_*i,t*_)(*z*_*i,t*_ *− θ*)]^*T*^, then the neuron solves:

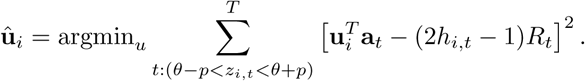

Performing stochastic gradient descent on this minimization problem gives the learning rule:

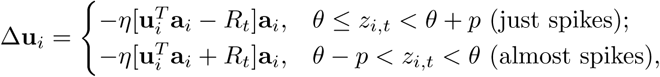

for all time periods at which *z*_*i,t*_ is within *p* of threshold *θ*.

### The causal effect as a finite-difference operator

The estimator can also be considered as a type of finite-difference approximation to the reward gradient we would compute in the deterministic case. This relation is fleshed out here. Specifically, we show that

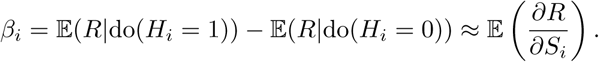

To show this we replace 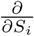 with a type of finite difference operator:

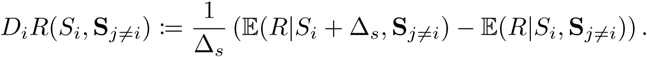

Here **S**_*j* ≠ *i*_ *⊂ 𝒳* is a set of nodes that satisfy the back-door criterion with respect to *H*_*i*_ → *R*. When *R* is a deterministic, differentiable function of **S** and Δ_*s*_ → 0 this recovers the reward gradient 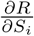 and we recover gradient descent-based learning.

To consider the effect of a single spike, note that unit *i* spiking will cause a jump in *S*_*i*_ compared to not spiking (according to synaptic dynamics). If we let Δ_*s*_ equal this jump then it can be shown that 𝔼(*D*_*i*_*R*) is related to the causal effect.

First, assuming the conditional independence of *R* from *H*_*i*_ given *S*_*i*_ and **S**_*j*≠*i*_:

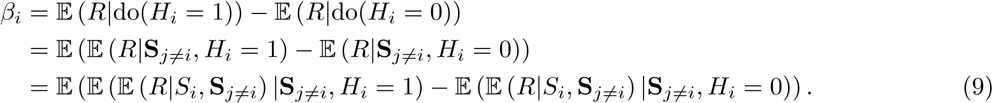

Now if we assume that on average *H*_*i*_ spiking induces a change of Δ_*s*_ in *S*_*i*_ within the same time period, compared with not spiking, then:

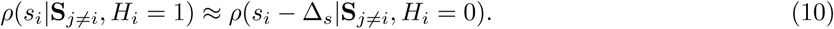

This is reasonable because the linearity of the synaptic dynamics means that the difference in *S*_*i*_ between spiking and non-spiking windows is simply exp(− *t*_*si*_*/τ*_*s*_)*/τ*_*s*_, for spike time *t*_*si*_. We approximate this term with its mean:

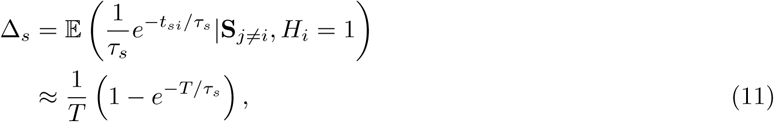

under the assumption that spike times occur uniformly throughout the length *T* window. These assumptions are supported numerically (Figure 6).

**Figure 6:**
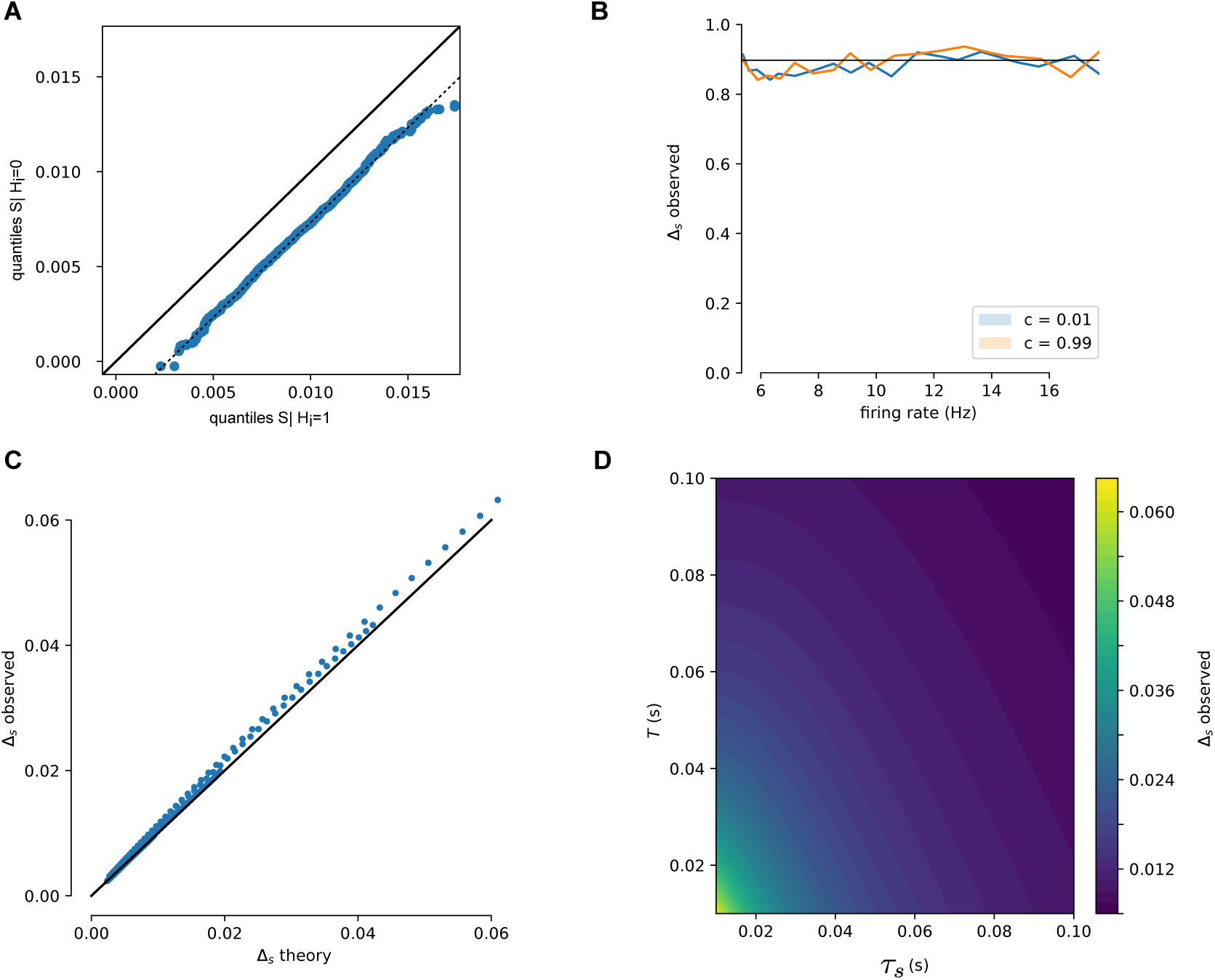
Relation between *S*_*i*_ and *H*_*i*_ over window *T*. (A) Simulated spike trains are used to generate *S*_*i*_|*H*_*i*_ = 0 and *S*_*i*_|*H*_*i*_ = 1. QQ-plot shows that *S*_*i*_ following a spike is distributed as a translation of *S*_*i*_ in windows with no spike, as assumed in (10). (B) This offset, Δ_*s*_, is independent of firing rate and is unaffected by correlated spike trains. (C) Over a range of values (0.01 *< T <* 0.1, 0.01 *< τ*_*s*_ *<* 0.1) the derived estimate of Δ_*s*_ (Equation (11)) is compared to simulated Δ_*s*_. Proximity to the diagonal line (black curve) shows these match. (D) Δ_*s*_ as a function of window size *T* and synaptic time constant *τ*_*s*_. Larger time windows and longer time constants lower the change in *S*_*i*_ due to a single spike.

**Figure 7:**
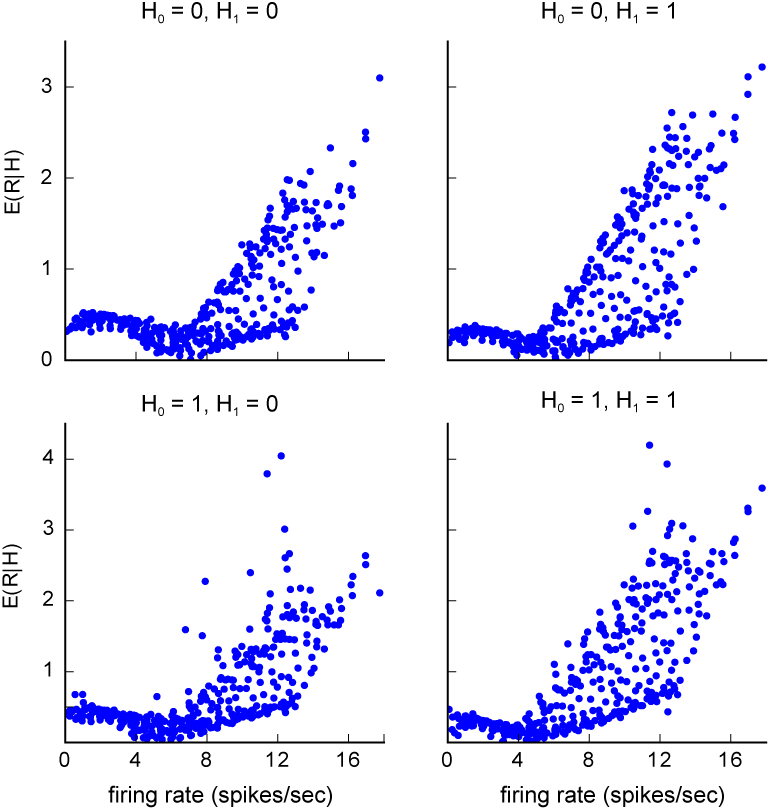
Supplementary Figure 1. **Validity of Assumption 1 for LIF network model.** Independence of 𝔼(*R*|*H*) from **w**. Over a range of network weights, the expected reward is shown here for all possible values of *H*_0_, *H*_1_. For low firing rates, the expected reward does not change much, regardless of network weight, showing the assumption is valid. For high firing rates this is not the case.

**Figure 8:**
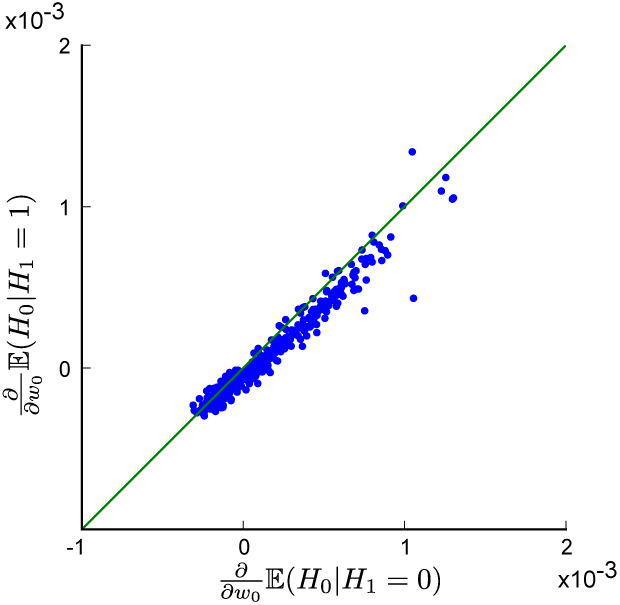
Supplementary Figure 2. **Validity of Assumption 2 for LIF network model.** Independence of 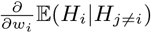 from *H*_1_. Over a range of network weights, the gradient 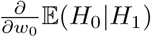 are roughly equal, regardless of the value of *H*_1_.

**Figure 9:**
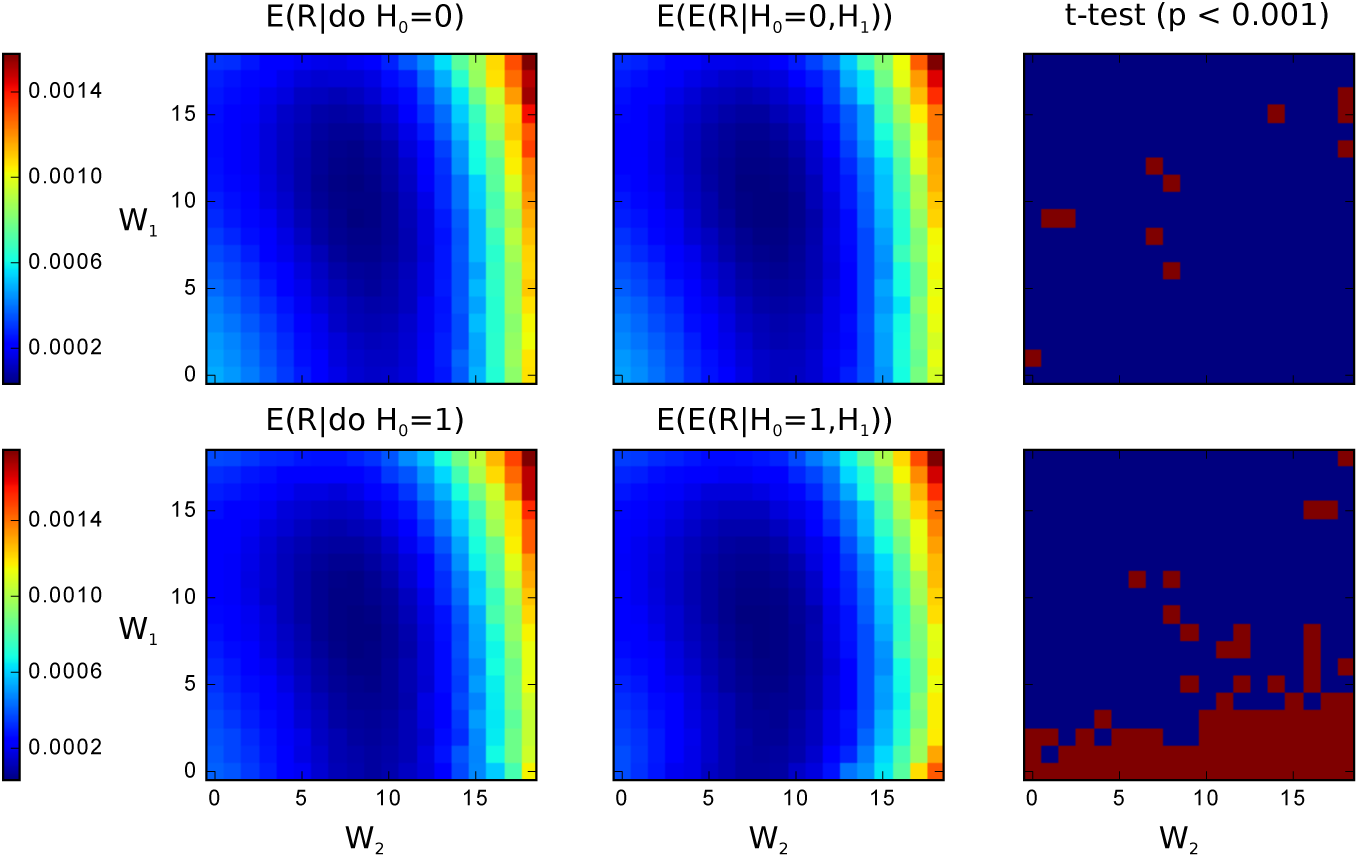
Supplementary Figure 3. **Validity of Assumption 3 for LIF network model.** Verifying the interventional distribution of expected reward (left) matches the backdoor corrected expected reward from observational data (middle). Samples from each distribution are compared via t-test (*α* = 0.001, Bonferroni corrected), to see if they have significantly different means (right; red = significant). The majority of network weights result in activity in which the interventional distribution matches the corrected observational distribution.

Writing out the inner two expectations of (9) gives:

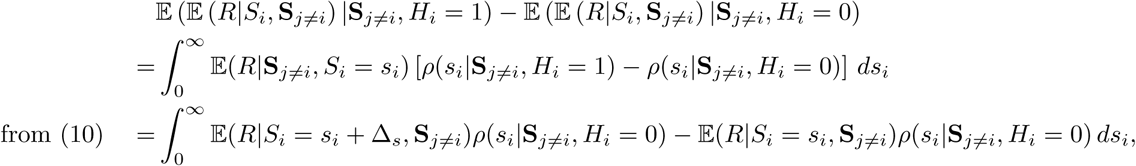

after making the substitution *s*_*i*_ → *s*_*i*_ + Δ_*s*_ in the first term. Writing this back in terms of expectations gives:

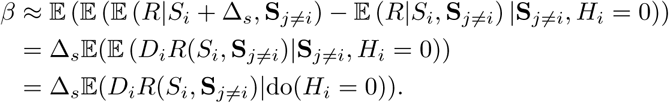

Thus estimating the causal effect is similar to taking a finite difference approximation of the reward gradient.

### Implementation

`python` code used to run simulations and generate figures is available at https://github.com/benlansdell/rdd.

## Acknowledgements

The authors would like to thank Roozbeh Farhoodi, Ari Benjamin and David Rolnick for valuable discussion and feedback.

## Author contributions

K.P.K and B.J.L. devised the study, B.J.L. performed the analysis, and K.P.K and B.J.L. wrote the manuscript.

## Supplementary material

### 1 Validity of causal assumptions for a leaky integrate-and-fire (LIF) network

The relation derived in the Methods between causal effect and policy gradient methods relied on three assumptions:

1. 𝔼 (*R*|*H*) is independent of *w*,
2. 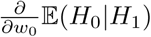 is independent of *H*_*j*≠*i*_,
3. *H*_*j*≠*i*_ satisfies the backdoor criterion with respect to *H*_*i*_ → *R*.

These assumptions are reasonable for certain parameter regimes of the LIF network used in this analysis (Supplementary Figure 1,2,3).

